# The Association between Cigarette Affordability and Consumption

**DOI:** 10.1101/361287

**Authors:** Yanyun He, Ce Shang, Frank J. Chaloupka

## Abstract

**Objectives:** To calculate cigarette affordability for 78 countries worldwide from 2001 to 2014 using the Relative Income Price (RIP) ratio defined as the percentage of per capita GDP required to purchase 100 packs of cigarettes using the lowest price from Economist Intelligence Unit (EIU) database, examine the association between cigarette affordability and cigarette consumption, and calculate the affordability elasticity of demand.

**Design and Methods:** RIP (2001-2014) was calculated for 16 low-income economies, 19 lower middle-income economies, 13 upper middle-income economies, and 30 high-income economies. Ordinary least square regressions were used to analyze the association between cigarette affordability and consumption.

**Results and Conclusions:** Per capita consumption continued to rise in low-income countries and decreased slightly in lower middle-income countries as the RIP of cigarette consistently declined in low- and lower middle-income economies from 2001 to 2014. The real cigarette prices continued to decline in low- and lower middle-income countries and continued to rise in upper middle- and high-income countries. Though cigarettes were more expensive in HICs than were in LMICs, cigarettes were more affordable in HICs than were in LMICs. The regression results show a 10% increase in the RIP of cigarettes led to a 2% decrease in per capita consumption. The affordability elasticity of demand differed significantly between HICs and LMICs. However, the effect of cigarette affordability on consumption has not changed over time. To control the smoking epidemic, low- and lower middle-income countries should further increase cigarette prices. The rate of price increase should exceed the rate of economic growth and outpace the inflation rate to make cigarettes less affordable and thereby reducing tobacco use.

## 1. Introduction

Numerous studies have shown that an increase in cigarette prices decreases cigarette consumption,(1-3) decreases smoking initiation, and increases quitting.(4-8) The main goal of increasing cigarette taxes, which results in higher cigarette prices, is to make cigarettes less affordable and thereby reducing the demand for cigarettes. Cigarette affordability, rather than the price of cigarettes alone, is a key determinant of the demand for cigarettes because the increase in income may erode the effects of taxes or prices by making cigarettes more affordable. This has happened to many low- and middle-income countries (LMICs) since 1990s and marked a major tobacco control failure.(9, 10) Therefore, examining the effect of cigarette affordability, rather than just price, on cigarette consumption can provide us more insights into the demand for cigarettes. In recent years, the tobacco control community has increasingly recognized tobacco affordability as a key determinant of tobacco use behavior in that it combines the impact of changing tobacco prices and economic growth. The analysis of the trend in cigarette affordability in a specific country is of great importance as it can advise policy makers whether the cigarette affordability has caught up with the pace of economic growth and whether there is room for further increase in cigarette prices through taxation. The cross-country comparison can also inform policy makers how cigarette affordability in a specific country compared to other countries in the world.

A number of definitions have been developed recently, such as Relative Income Price (RIP) and the minutes of labor required to purchase a pack of cigarettes.(9-11) RIP, developed by Blecher and Van Walbeek,(9) is defined as the percentage of per capita GDP required to purchase 100 packs of cigarettes. The higher the RIP is, the less affordable cigarettes become and vice versa. Furthermore, the RIP ratio is a relative measure that allows for comparisons over time and across countries, and thus has advantages in capturing and comparing the real cost of cigarettes. Blecher and Van Walbeek (2009) has shown that RIP is most appropriate when measuring affordability in LMICs.(10) For high-income countries (HICs), the choice of income measure is not important. Therefore, RIP will be used as the measure of affordability in this present study.

Both economic theory and empirical evidence clearly underscore the importance of affordability as a determinant of tobacco use as consumers make decisions based on income and price simultaneously. In other words, affordability mediates the effects of tax/price and income on tobacco use. Kostova et al. (2015) examined smoking initiation and cessation in LMICs and found that price effects in promoting cessation and in preventing initiation differed by income levels, suggesting that cigarette affordability, rather than prices, played a major role in smoking behavior.(12) A limited number of studies have explicitly examined cigarette affordability by using descriptive analysis. Guindon et al. (2002) compared cigarette price data from more than 80 countries using varying methods and examined trends in prices and affordability during the 1990s. (11) Cigarette affordability was defined as the minutes of labor required to purchase a pack of cigarettes in their study. The prices of cigarettes were presented in US dollars, purchasing power parity (PPP) units using the Big Mac index as an indicator of PPP. The authors found that cigarette prices tended to be higher in wealthier countries and in countries that had strong tobacco control programs. Trends between 1990 and 2000 in real prices and minutes of labor indicated, with some exceptions, that cigarettes had become more expensive in most developed countries but more affordable in many developing countries. Cigarette prices have failed to keep up with increases in the general price level of goods and services, rendering them more affordable in 2000 than they were at the beginning of the decade. Kan (2007) used another definition of affordability, the ratio of the price of one pack of cigarettes to daily income and compared the cigarette affordability in 60 cities worldwide in 2006.(13) The author found that cigarette affordability in most of the surveyed cities remained high, which suggested that there was room for increasing cigarette prices via tax increases. The author also pointed out that the increase in cigarette prices in newly emerging economies, such as China, India, and Singapore, has lagged behind the high speed of economic growth and opportunities to increase taxes and government revenues have been overlooked. Blecher & Van Walbeek (2009) calculated cigarette affordability for a number of countries using two different definitions of affordability, which were Relative Income Price (RIP) and minutes of labor required to purchase the cheapest pack of cigarettes, based on net earnings to investigate trends between 1990 and 2006.(10) They also assessed the appropriateness of different measures of affordability. They found that in HICs cigarettes were significantly more affordable than in LMICs, but had become less affordable since 1990. Among LMICs, cigarettes had become more affordable since 1990 and at an increasingly rapid rate since 2000. Similarly, several other studies examined the trend in cigarette affordability in different parts of the world for different time periods.(14-16) Blecher & Van Walbeek (2004) was the only study so far to investigate the relation between cigarette affordability and consumption by estimating the affordability elasticities of demand.(9) Seventy countries were investigated, of which 28 and 42 were categorized as developed and developing countries, respectively. The affordability of cigarettes was defined as the percentage of per capita GDP required to purchase 100 packs of cigarettes (RIP). They found that a 1% increase in the RIP was expected to decrease cigarette consumption by between 0.49 - 0.57%. They also found that the affordability elasticity of demand did not differ significantly between rich and poor countries as did the price elasticity of demand.

To sum up, among the aforementioned studies only Blecher and Van Walbeek (2004) examined the association between cigarette affordability and consumption; all other studies were simply descriptive analysis of cigarette affordability for different countries during different time periods. Whether the conclusions of Blecher & Van Walbeek (2004) will still be valid nowadays is unknown. Extending on the work of Blecher & Van Walbeek (2004), this study aims to investigate the association between cigarette affordability and consumption using data from 2001 to 2014 to see if using data from recent years will still support their findings. The latest statistics and trends will be examined and presented as well.

## 2. Methods

### 2.1. Data and Measures

#### World Bank World Development Indicator (WDI) database

WDI database is time series data compiled from officially recognized international sources. Aggregated country level characteristics from 2001-2014, such as per capita GDP, percentage of female population, unemployment rate, percentage of population aged 15-64, and country classification are directly drawn from the WDI database. Country classification is defined according to per capita Gross National Income (GNI) in US dollars using the World Bank Atlas method. 78 countries are classified into low-income countries (≤$745), lower middle-income countries ($746-$2,975), upper middle-income countries ($2,976-$9,205), and high-income countries (>$9,205) by the year of 2001 standard. Specifically, low-income, lower middle-income, upper middle-income, and high-income economies consisted of 16, 19, 13, and 30 countries, respectively. The detailed country list of each category was shown in the Appendix. The information on Purchasing Power Parity (PPP) conversion factor and Consumer Price Index (CPI) were also sourced from the WDI database.

#### Economist Intelligence Unit (EIU)

The EIU city data collects cigarette prices for Marlboro or other leading international brands and a local brand sold at supermarkets and mid-priced stores in 140 cities in 92 countries from 2001 to 2014. Since this study focuses on cigarette affordability, the lowest of the four prices (two brands in two types of retail outlets) was selected to conduct the analysis. Cigarette prices and per capita GDP in local currency were used when constructing the affordability measure to ensure its accuracy. However, the nominal and real PPP adjusted price were used in cross-country comparison.

#### EuroMonitor International

Two measures of cigarette consumption were used in the analysis, -consumption that was realized at retail outlets and all cigarette consumption that included illicit trade. The per capita cigarette consumption in sticks in each country and year was derived using the aggregated country level cigarette consumption divided by the population aged 15 and over provided by the WDI database. In the regression analysis, the affordability measure and per capita cigarette consumption were transformed into log forms. By making this transformation, the estimated coefficient of affordability would represent the affordability elasticity of demand directly.

#### MPOWER Scores

The MPOWER measures were introduced by the WHO under the Framework Convention on Tobacco Control (FCTC) guidelines to assist countries in achieving their tobacco control goals using six known tobacco control methods: M (monitor tobacco use), P (protect people from smoke), O (offer help to quit), W (warn about the dangers of tobacco), E (enforce bans on tobacco marketing), and R (raise taxes on tobacco). The database contains scores that evaluate these six policy dimensions in 196 countries in 2007-2008, 2010, 2012 and 2014. Six MPOWER scores were categorized into four or five levels. For the M policy dimension, the score value ranges from 1 to 4 in which a score of 1 represents no known data or no recent data (since 2009) or data that is neither recent nor representative, and a score of 2-4 represents the strength of policy from the weakest to the strongest.(17) For the other 5 policy dimensions (POWER), the scores measure overall strength on a scale of 1 to 5 in which a score of 1 represents a lack of data (missing data) and a score of 2-5 represents the weakest to the strongest policies. Since the MPOWER scores were not reported in years 2009, 2011, and 2013, we assume that there were no policy changes over those three years and approximate the scores by using the scores in the prior years (2008, 2010, and 2012). Then a composite score was constructed for each country and year by summing up each individual scores.

### 2.2. Methodology

Equation (1) was used to examine the association between per capita cigarette consumption and cigarette affordability.

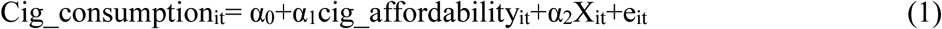

i denotes country and t denotes year. In specification (1), covariates include percentage of population aged 15-64, percentage of female population, and unemployment rate. Specification (2) includes all the covariates used in specification (1) plus the MPOWER composite score, which is the summation of the six MPOWER policy variables. Since the MPOWER composite scores are only available from 2007 onward, specification (1) was used for the sample from 2001 to 2014, whereas specification (2) was used for the sample from 2007 to 2014. The lowest prices from the EIU database were used to construct the affordability measure for specification (1), whereas the prices for the most sold brand from MPOWER package were used in specification (2) for comparison. Variance Inflation Factor (VIF) was estimated to evaluate whether the country-level affordability measure were highly correlated with time-invariant country-level unobservable attributes. The Ordinary Least Square (OLS) technique was used to carry out the analysis for each specification. The standard errors were clustered at the country level to account for the correlations between years.

## 3. Results

### 3.1 Trend in Affordability from 2001 to 2014

The trend in affordability using the EIU lowest prices from 2001 to 2014 is shown in Table 1 and Figure 1. Only countries with non-missing data for all years are included in the analysis to reflect the trend during the entire period. This results in 78 countries in the analysis. For low-and lower middle-income countries, the overall trend in the RIP of cigarettes is decreasing, indicating that cigarettes have become more affordable since 2001. In upper middle- and high-income countries cigarettes have become less affordable over time since 2001. The percentage change from 2001 to 2014 in the RIP is -3.16%, -32.79%, 138.84%, and 63.15% for low-, lower middle-, upper middle-, and high-income countries, respectively. Comparing the RIP of cigarettes across all country categories, cigarettes were more affordable in high-income countries than were in low-income countries.

**Table 1:** Relative Income Price of cigarettes at EIU lowest prices 2001-2014

| Year | Low-Income | Lower Middle-Income | Upper Middle-Income | High-Income |
| --- | --- | --- | --- | --- |
| 2001 | 0.259 | 0.167 | 0.022 | 0.010 |
| 2002 | 0.327 | 0.163 | 0.022 | 0.010 |
| 2003 | 0.327 | 0.161 | 0.029 | 0.010 |
| 2004 | 0.295 | 0.196 | 0.030 | 0.011 |
| 2005 | 0.259 | 0.186 | 0.030 | 0.011 |
| 2006 | 0.234 | 0.128 | 0.027 | 0.011 |
| 2007 | 0.256 | 0.157 | 0.028 | 0.011 |
| 2008 | 0.252 | 0.116 | 0.030 | 0.012 |
| 2009 | 0.288 | 0.108 | 0.033 | 0.013 |
| 2010 | 0.230 | 0.105 | 0.034 | 0.013 |
| 2011 | 0.210 | 0.101 | 0.037 | 0.014 |
| 2012 | 0.224 | 0.104 | 0.044 | 0.015 |
| 2013 | 0.243 | 0.120 | 0.046 | 0.015 |
| 2014 | 0.250 | 0.112 | 0.052 | 0.016 |

Figure 1-Average Annual Percentage Change in the RIP, 2001-2014

Figure 2 plots the average annual percentage change in the RIP from 2001 to 2014 at the country level. There are 79 countries in this plot because Libya has the affordability data in the year of 2001 or 2014, but had missing data in other years, which resulted being excluded from Figure 1. Cigarettes have become more affordable in 12 countries, -4 low-income countries (Azerbaijan, Cambodia, Nigeria, and Uzbekistan), 6 lower middle-income countries (Algeria, China, Colombia, Morocco, Paraguay, Peru, and Tunisia), and 2 high-income countries (Hong Kong, China and Qatar). Two biggest increases in RIP took place in Argentina and Serbia, with average annual percentage change being 38.98% and 34.01%, respectively.

Figure 2-Average Annual Percentage Change in the RIP, 2001-2014

Figure 3 and 4 show the trend in nominal and real cigarette prices from 2001 to 2014. First, cigarette prices were converted into a common dollar currency using the Purchasing-Power Parity (PPP) conversion factor. The PPP factor is the number of local currency units required to buy the same amount of goods or services in the domestic market as US $1 in the USA. The nominal PPP adjusted cigarette prices in each country and year were further adjusted for inflation using the CPI (base year 2010). To reflect the true trend in nominal/real cigarette prices, only countries with non-missing nominal/real price data for all years were included in the analysis. This results in 78 countries in the analysis of nominal price and 72 countries in the analysis of real price (See Appendix 1). Six countries were excluded from the real price analysis due to the missing of CPI data. The trend in nominal cigarette prices is pretty flat for low- and lower middle-income countries. Nominal cigarette prices have been decreasing since 2001 for low-income countries, and slightly increasing for lower middle-income countries. However, the nominal cigarette prices have been consistently increasing in upper middle- and high-income countries. The percentage change in nominal cigarette prices from 2001 to 2014 is -8.33%, 7.65%, 91.44%, and 83.98% for low-, lower middle-, upper middle-, and high-income countries, respectively. Figure 4 shows the trend in inflation adjusted cigarette prices in PPP. Since many low- and lower middle-income countries underwent fast growing in economy during the past decade and high inflation usually accompanied the fast economic development, real cigarette prices exhibit a distorted trend, especially for the time period before 2009 when the real cigarette prices in low- and lower middle-income countries were even higher than were in upper middle- and high-income countries. The real cigarette prices in low- and lower middle-income countries have been consistently decreasing due to the fact that the increase in cigarette prices failed to catch up with inflation. The percentage change in real cigarette prices from 2001 to 2014 is - 57.57%, -52.81%, 14.53%, and 39.24% for low-, lower middle-, upper middle-, and high-income countries, respectively. Figure 3 and 4 indicate that though cigarettes were more affordable in HICs, the real prices for cigarettes remained higher than LMICs by year 2014.

Figure 3-Nominal PPP adjusted EIU lowest price

Figure 4-Real PPP adjusted EIU lowest Price in 2010 dollars

The trend in cigarette per capita retail consumption is shown in Figure 5. For high- and upper middle-income countries, cigarette per capita consumption has been consistently decreasing since 2001. However, on the other hand, cigarette per capita consumption has been gradually increasing since 2001 in lower middle-income countries until 2009 and low-income countries until 2007, then entered into a downward trend afterwards. The percentage change in per capita consumption from 2001 to 2014 is 16.87%, -15.85%, -32.69%, and -39.04% for low-, lower middle-, upper middle-, and high-income countries, respectively. The trend in cigarette per capita consumption echoed with the trend in RIP of cigarettes (see Figure 1), demonstrating a negative association between the two. It is worth noting that per capita consumption has been higher in lower middle-income countries than in upper middle-income countries since 2004 and drawn very close to that in high-income economies since 2009 despite the fact that the RIP of cigarettes was much higher in lower middle-income countries during most of the analysis period.

Figure 5-Cigarette Per Capita Consumption_Retail

### 3.2. Regression Results

Table 2 presents the summary statistics of the variables used in specification (1) and (2). The magnitudes of VIF are over 30 when regressing each affordability measure on country fixed effects, indicating that high multicollinearity exists between each affordability measure and the country-specific attributes. However, the VIFs are only around 1 when regressing each affordability measure on year fixed effects, suggesting that no multicollinearity exists between each affordability measure and the year-specific attributes. It also indicates that the variations of the affordability measures are mainly within year and between country variations. Thus, OLS with year fixed effects was also estimated for the purpose of comparison.

**Table 2:** Summary statistics of the variables used in the regression.

| Variables | Specification (1) | Specification (2) |
| --- | --- | --- |
| Per capita cigarette retail consumption in sticks | 1298.185<br>(821.724) | 1288.609<br>(823.870) |
| Log of per capita cigarette consumption in sticks | 6.904<br>(.832) | 6.894<br>(.832) |
| Cigarette affordability 1 (EIU lowest price per pack/ GDP per Capita*100) | .107<br>(.367) |  |
| Log of cigarette affordability 1 | -3.501<br>(1.207) |  |
| Cigarette affordability 2 (MPOWER most popular price per pack/ GDP per Capita*100) |  | .106<br>(.336) |
| Log of cigarette affordability 2 |  | -3.284<br>(1.115) |
| Population aged 15-64, % of total | 66.168<br>(4.885) | 66.585<br>(4.637) |
| Population female, % of total | 50.175<br>(3.159) | 50.361<br>(3.245) |
| Unemployment, % of total labor force | 7.847<br>(4.586) | 8.501<br>(5.248) |
| Composite MPOWER Score |  | 20.778<br>(3.236) |
| N | 887<sup>1</sup> | 519<sup>2</sup> |

Note:
1. This sample contains 64 countries.
2. This sample contains 75 countries.

Table 3 presents the regression results of specification (1) and (2). The coefficients of the affordability measure were pretty robust regardless of what affordability measure was used and whether the composite MPOWER score and year fixed effects were controlled for. Roughly speaking, per capita cigarette consumption dropped by 2 % as the RIP of cigarettes went up by 10%. The estimates of the affordability measure were significant at 0.1% level, clearly indicating the strong association between per capita cigarette consumption and affordability. Per capita cigarette consumption increased as the percentage of population aged 16-64, the percentage of female population, and the unemployment rate increased. The effect of composite MPOWER score on per capita cigarette consumption was not significant. The regression results that examined the effect of cigarette affordability on per capita all consumption, which included illicit trade, were shown in Appendix 2.

**Table 3:**
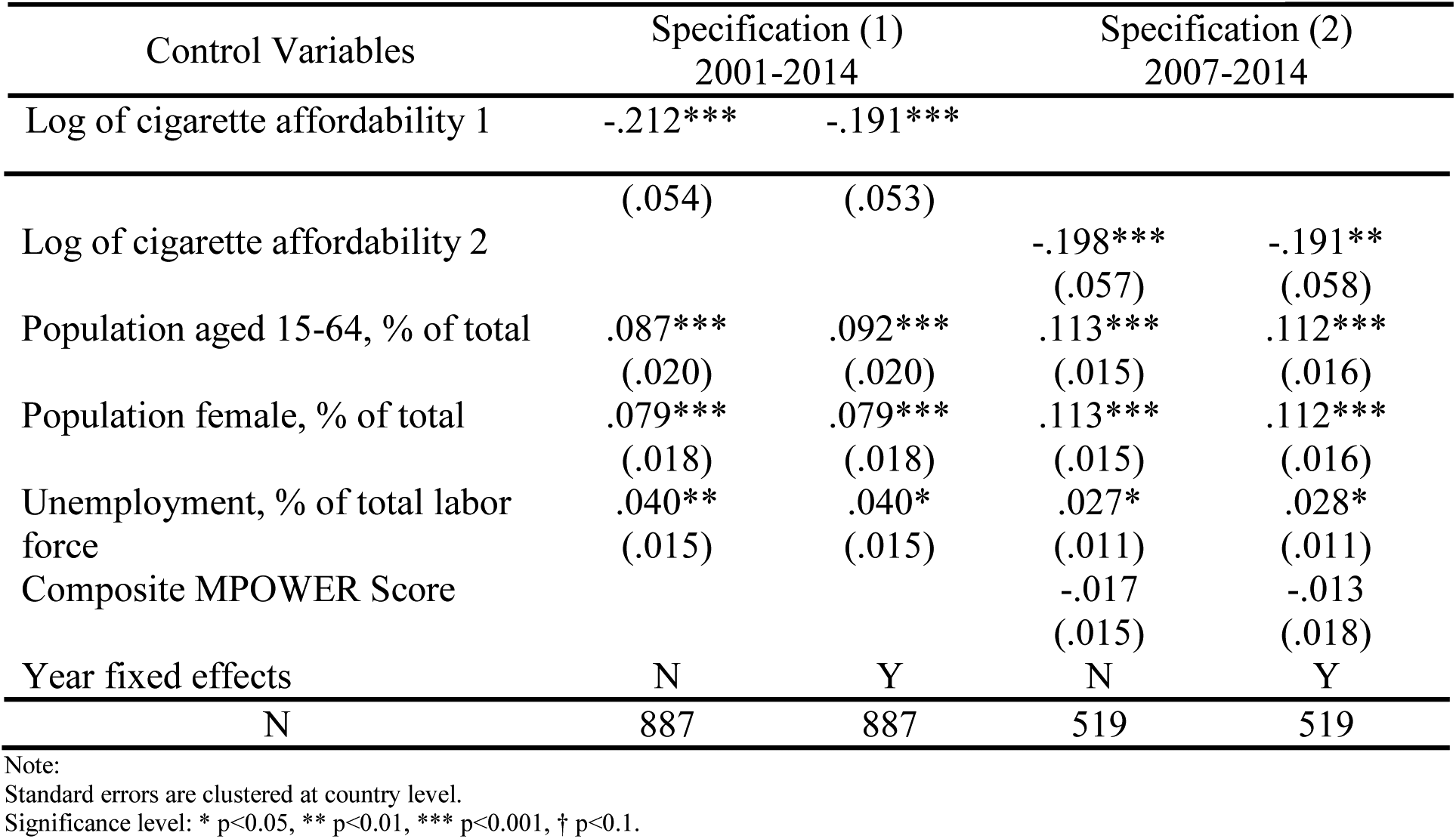
The effect of cigarette affordability on per capita retail consumption.

To see whether the affordability elasticity of demand will differ across country categories, we grouped all countries into two groups, -HICs and LMICs, and stratified the analysis of specification (1). The results are shown in Table 4. The affordability elasticity of demand was bigger in HICs than was in LMICs. Per capita cigarette consumption dropped by 10% if the RIP of cigarettes went up by 10% in HICs, whereas less than 2% in LMICs, indicating that cigarette affordability was more elastic in HICs than was in LMICs. A further full interaction test shows that the effect of cigarette affordability on per capita consumption differed significantly between HICs and LMICs (see Table 5). To see whether the affordability elasticity of demand has changed over time, we further implemented a full interaction test of year dummy and all control variables in specification (1) year fixed effects model. The results show that all the interaction terms of year dummy and affordability measure collectively were not significant, indicating that the effect of affordability on consumption did not change over time. The results will be available upon request.

**Table 4:** Stratified regression results of specification (1) by economy type.

| Control Variables | Overall | LMICs | HICs |
| --- | --- | --- | --- |
| Log of cigarette affordability at lowest prices | -.191***<br>(.053) | -.142*<br>(.058) | -.957***<br>(.211) |
| Population aged 15-64, % of total | .092***<br>(.020) | .116***<br>(.019) | -.025<br>(.042) |
| Population female, % of total | .079***<br>(.018) | .078<br>(.062) | .041<br>(.034) |
| Unemployment, % of total labor force | .040*<br>(.015) | .035†<br>(.018) | .045**<br>(.014) |
| Year fixed effects | Y | Y | Y |
| N | 887 | 565 | 322 |

Note: Standard errors are clustered at country level. Significance level: \* p<0.05, \*\* p<0.01, \*\*\* p<0.001, † p<0.1.

**Table 5:** Full interaction test of high income indicator and all other covariates.

| Control Variables | Coefficient |
| --- | --- |
| Log of cigarette affordability at lowest prices | -.148*<br>(.058) |
| High-income indicator | 8.604<br>(5.474) |
| Interaction of affordability and high-income | -.706***<br>(.172) |
| Year fixed effects | Y |
| N | 887 |

Note: Standard errors are clustered at country level. Significance level: \* p<0.05, \*\* p<0.01, \*\*\* p<0.001, † p<0.1.

## 4. Discussions

Though the effect of cigarette prices and income on cigarette use has been well examined, only limited number of literature have examined the effect of cigarette affordability, a combined measure of cigarette prices and per capita GDP, on cigarette consumption. This study extends on the work of Blecher & Van Walbeek (2004) by examining the association between cigarette affordability and consumption employing the aggregate country-level data from 2001 to 2014. This study has two limitations. First, various country-specific tobacco control measures, such as anti-smoking campaign and programs, smoking bans, advertising bans, and health warning labels, etc., can potentially confound the results in specification (1) for the 2001-2014 sample. However, the estimates of affordability did not change much from specification (1) to (2) which incorporated six MPOWER policy indices into the analysis for the 2007-2014 sample. Secondly, the estimates of the affordability measure in Table 4 are likely to be biased due to the omission of country level stage of smoking epidemic. As noted in tobacco control country profiles published by WHO in 2003,(18) “Stage 1 of the WHO paradigm is characterized by a low prevalence (below 20%) of cigarette smoking, principally limited to males. Stage 2 of the smoking epidemic is characterized by increases in smoking prevalence to above 50% in men, early increases in cigarette smoking among women. Stage 3 of the epidemic is characterized by a marked downturn in smoking prevalence among men, a more gradual decline in women, and convergence of male and female smoking prevalence. Stage 4 of the epidemic is characterized by a marked downturn in smoking prevalence in both men and women.” Since cigarette consumption rises as the stage of smoking epidemic goes up in countries of Stage 1, 2, and 3 and declines in countries of Stage 4,(9) and most LMICs were in Stage 1, 2, or 3 and HICs were in Stage 4, omitting country level stage of smoking epidemic would result in a downward bias of the affordability measure for HICs and upward bias for LMICs.

Two differences were found when comparing our results with Blecher & Van Walbeek (2004). One is concerning the estimate of the affordability elasticity of demand, which is -0.20 in our study and -0.49-0.57 in Blecher & Van Walbeek (2004). Two reasons may give rise to this difference. First, the data were from different time period, -1990-2001 in Blecher & Van Walbeek (2004) study and 2001-2014 in our study. Secondly, Blecher & Van Walbeek (2004) study was a bivariate regression, which did not include any other covariates in the analysis. Therefore, the estimates of the affordability elasticity in their study was likely to be biased. We tested the bivariate regression for specification (1) and (2) in our study and got around -0.30 for the affordability elasticity of demand. As we pointed out earlier, omitting country-level tobacco control policies in our analysis would bias the estimate of affordability downward for HICs and upward for LMICs. However, the overall direction of bias is hard to predict. Another difference is concerning whether the affordability elasticity of demand differed between rich and poor countries. Blecher and Van Walbeek (2004) found that the affordability elasticity of demand did not differ significantly between rich and poor countries as did the price elasticity of demand.(9) However, we found that the effect of cigarette affordability on per capita consumption differed significantly between HICs and LMICs. Potential downward bias of the affordability elasticity for HICs and upward bias for LMICs in our analysis may contribute to this discrepancy.

## 5. Conclusions

In this paper we employed the OLS technique to examine the affordability elasticity of cigarette demand from 2001 to 2014 for 64 countries worldwide. The trend in cigarette affordability, nominal/real cigarette prices, and per capita cigarette consumption were also explored and presented. We further stratified the analysis by country category. The results clearly show that per capita cigarette consumption was negatively associated with cigarette affordability, indicating that smokers consumed less cigarettes as cigarettes became less affordable. The affordability elasticity of demand differed significantly between HICs and LMICs. However, the effect of cigarette affordability on consumption had not changed over time. Cigarettes have been more affordable in low- and lower middle-income economies and less affordable in upper middle- and high-income countries during the study period. The real cigarette prices continued to decline in low- and lower middle-income countries and continued to rise in upper middle- and high-income countries. Echoing with the dramatic decrease of real cigarette prices in low-income countries, the per capita cigarette consumption has been increasing since 2001 in these countries, resulted in serious challenges in tobacco control. To control the smoking epidemic, low- and lower middle-income countries should further increase cigarette prices. The rate of increase in cigarette prices should exceed the rate of economic growth and outpace the inflation rate to make cigarettes less affordable and thereby reducing cigarette use.

**Appendix 1:** List of countries of each economy type included in Figure 1, 3, and 5

| Low-income economies | Lower middle-income economies | Upper middle-income economies | High-income economies |
| --- | --- | --- | --- |
| Azerbaijan† | Algeria | Argentina† | Australia* |
| Bangladesh* | China† | Brazil | Austria |
| Cambodia* | Colombia | Chile† | Bahrain* |
| Cameroon | Ecuador | Costa Rica | Belgium |
| Cote d'Ivoire* | Egypt, Arab Rep. | Czech Republic | Canada |
| India | Guatemala | Hungary | Denmark |
| Indonesia | Iran, Islamic Rep. | Malaysia | Finland |
| Kenya | Morocco | Mexico | France |
| Nepal* | Paraguay* | Oman* | Germany |
| Nigeria* | Peru | Panama* | Greece |
| Pakistan | Philippines | Poland | Hong Kong SAR, China |
| Papua New Guinea* | Romania | Saudi Arabia | Iceland* |
| Senegal* | Russian Federation | Uruguay | Ireland |
| Ukraine | Serbia |  | Italy |
| Uzbekistan† | South Africa |  | Japan |
| Vietnam | Sri Lank* |  | Korea, Rep. |
|  | Thailand |  | Kuwait* |
|  | Tunisia |  | Luxembourg* |
|  | Turkey |  | Netherlands |
|  |  |  | New Zealand* |
|  |  |  | Norway |
|  |  |  | Portugal |
|  |  |  | Qatar |
|  |  |  | Singapore |
|  |  |  | Spain |
|  |  |  | Sweden |
|  |  |  | Switzerland |
|  |  |  | United Arab Emirates† |
|  |  |  | United Kingdom |
|  |  |  | United States |
| 16 | 19 | 13 | 30 |

Note:
† denotes countries that were excluded from Figure 4 due to the missing of CPI
\* denotes countries that were excluded from regression analysis due to the missing of consumption data.

**Appendix 2:**
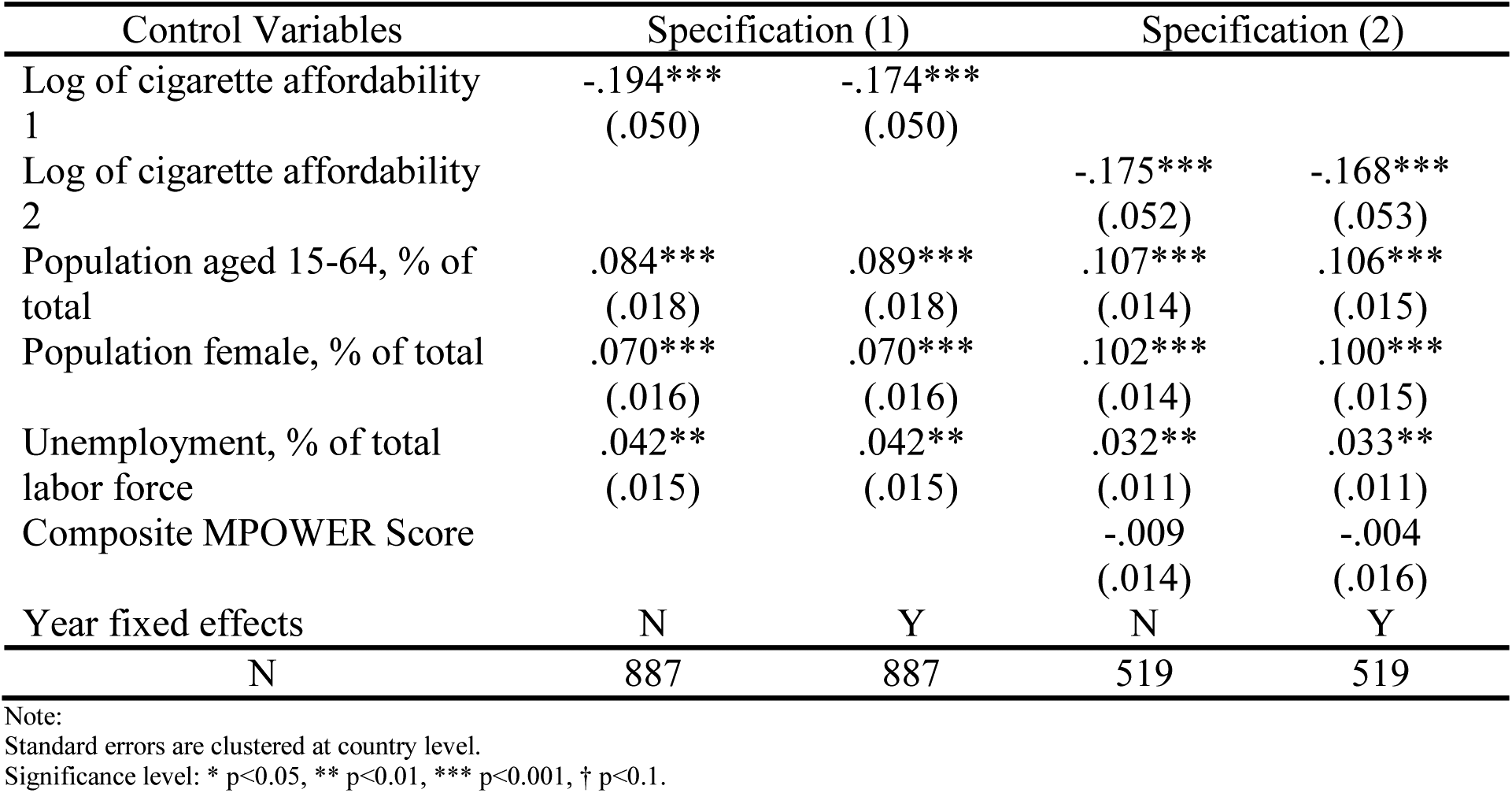
The effect of cigarette affordability on per capita ALL consumption

